# Se-Glargine I. Chemical Synthesis of a Basal Insulin Analog Stabilized by an Internal Diselenide Bridge

**DOI:** 10.1101/2023.06.23.546304

**Authors:** Orit Weil-Ktorza, Balamurugan Dhayalan, Michael A. Weiss, Norman Metanis

**Affiliations:** The Institute of Chemistry, The Hebrew University of Jerusalem Edmond J. Safra, Givat Ram, Jerusalem 9190401, Israel; Department of Biochemistry and Molecular Biology, Indiana University School of Medicine, Indianapolis, IN 46202 USA

**Keywords:** protein engineering, protein design, hormone, metabolism, diabetes mellitus

## Abstract

Insulin, a small globular protein, has long provided a model for studies of biophysical principles with therapeutic application. The safety and efficacy of insulin replacement therapy for the treatment for diabetes mellitus have been enhanced by protein engineering. Here, we describe the chemical synthesis of a basal insulin analog stabilized by the substitution of an internal cystine (A6-A11) by a diselenide bridge. The studies focused on insulin glargine, the active component of clinical products Lantus^®^ and Toujeo^®^ (Sanofi). Formulated in solution at pH 4 in the presence of zinc ions, insulin glargine exhibits a shifted isoelectric point (from pH 4.5 to neutral pH) due to a basic extension of the B chain (Arg^B31^-Arg^B32^). Subcutaneous injection of such an acidic formulation leads to pH-dependent precipitation of protein-zinc complexes to form a long-lived depot. Pairwise substitution of Cys^A6^ and Cys^A11^ by selenocysteine (Sec; the 21^st^ encoded amino acid) was effected by solid-phase peptide synthesis. The modified A chain also contained substitution of Asn^A21^ by Gly, introduced in glargine to avoid acid-catalyzed deamidation of the A21 carboxamide group in the formulation. Although classical chain combination of the di-Arg-extended B chain and modified A chain exhibited lower yield than does wild-type chain combination, substantial product was obtained through repeated reactions and successive purification. This strategy exemplifies the rational optimization of protein stability and may be generalizable to diverse disulfide-stabilized proteins of therapeutic interest.

---

Insulin provides a model for foundational studies of protein biophysics with potential therapeutic application (Fig. 1; ref 1). In this and the following article (2), we demonstrate that substitution of an internal cystine by a diselenide bridge in a basal insulin analog (3) provides a structure-based strategy to augment protein stability (4,5).^1^ The present study (Part I) focuses on the chemical synthesis (6) of an analog of insulin glargine in which cystine A6-A11 is substituted by a diselenide bridge (4). In the companion study (part II; ref 2) we report that native function (as probed in a cell-based model; 7) and structure (as probed by high-resolution NMR spectroscopy; 8) are maintained. The diselenide-stabilized analog, designated *Se-glargine*, may be of translational interest for the treatment of diabetes mellitus (DM) in the developing world (9).

**Figure 1.**
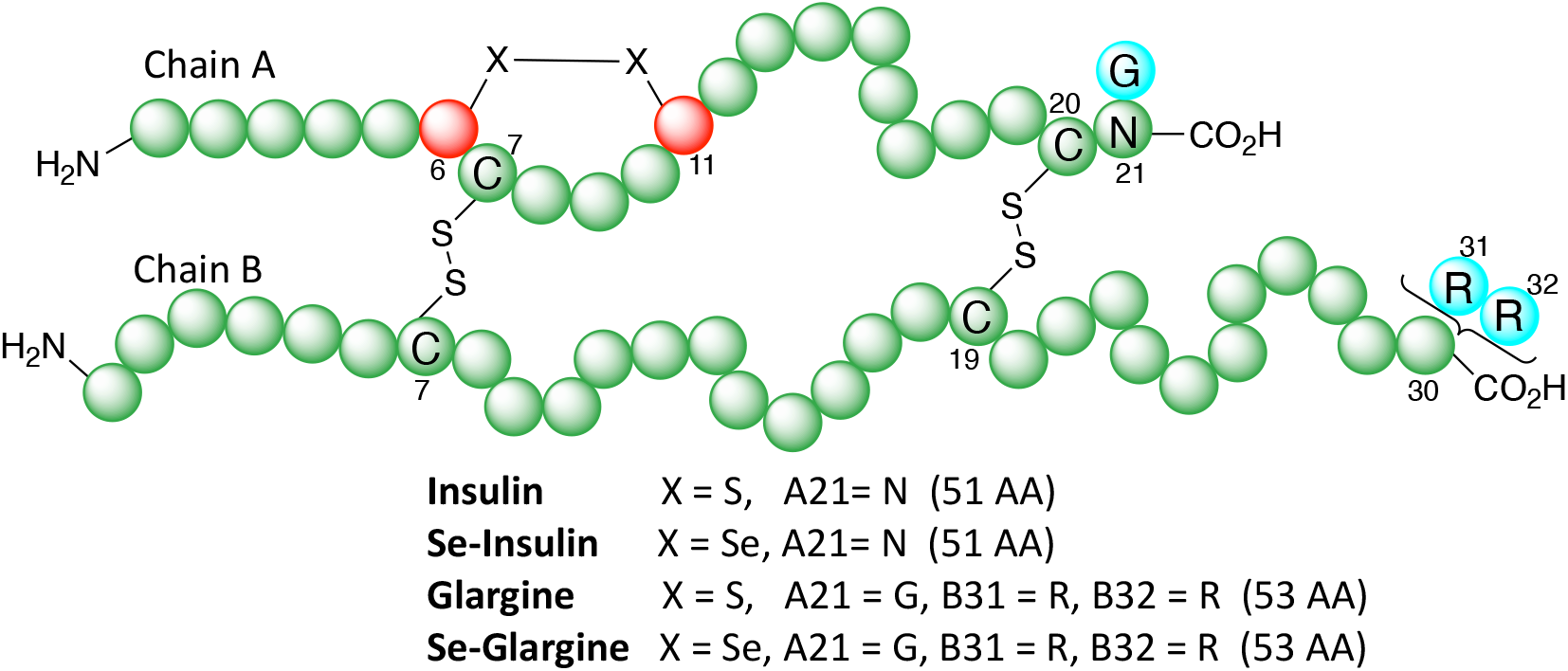
Schematic representation of the structure of insulin and glargine. The changes in glargine are shown in *cyan*. For Se-insulin and Se-glargine, the Cys residues at A6 and A11 (in *red*) were replaced by Sec.

Selenocysteine (Sec, U), a near isostere of Cys and the 21^st^ proteinogenic amino acid (10), has provided an elegant tool in peptide chemistry (11,12) and for enhancing the efficiency of protein folding (13-16). Because the selenol group of Sec has a lower p*K*_a_ (near 5.2; ref 17) and lower reduction potential (*E*_0_= -388 mV; ref 18) relative to the thiol group of Cys^2^, pairwise Cys-to-Sec substitutions can enhance the rate and efficiency of oxidative folding (13-16,19-21). Although such properties are pertinent to the nascent folding of proteins (including insulin; 4), the present study instead exploits geometric aspects of selenium: its larger atomic size and longer bond lengths relative to sulfur (22). We hypothesized that such steric features (unrelated to redox chemistry) could mitigate cryptic packing defects in the hydrophobic core of a globular protein (23) and so in favorable cases enhance stability (24). Proof of principle was provided in studies of wild-type insulin (4).

Application of diselenide chemistry to insulin glargine was motivated by its importance in global health (25,26) in the face of a growing pandemic of Type 2 diabetes mellitus (T2D) (27,28). Its formulation in an acidic solution nonetheless limits its shelf life at or above room temperature, as under these conditions insulin cannot form protective zinc-stabilized hexamers (29). We envision that Se-glargine will facilitate insulin replacement therapy in challenges regions lacking access to electricity or refrigeration (30). Although prepared herein by chemical synthesis and chain combination (4), advances in methods for recombinant Sec incorporation suggest that our strategy will be compatible with large-scale manufacture and generalizable to diverse proteins of therapeutic or industrial interest (31,32).

## Results

We previously described the preparation of human Se-insulin, in which the internal intrachain disulfide bond A6-A11 was replaced with diselenide via pairwise substitution of Cys^A6^ and Cys^A11^ by Sec (4). To this end, a modified 21-residue peptide, designated *A chain[C6U, C11U]*, was prepared in high yield by solid-phase peptide synthesis (SPPS); the wild-type B chain was prepared by sulfitolysis of human insulin (of either synthetic or biosynthetic origin). The purified chains were then mixed in the combination buffer to give, after only 8 h, the desired Se-insulin in 34% isolated yield (4). Whereas structure and activity were almost identical to wild-type (WT) insulin, the diselenide substitution enhanced stability by *ca*. 1 kcal/mol, delaying protease degradation and reductive unfolding (4).

Based on these results, we sought to test whether an analogous A6-A11 diselenide substitution could likewise stabilize a clinical insulin analog without affecting its structure and function. Insulin glargine was chosen based on the medical importance of products Lantus^®^ and Toujeo^®^ (Sanofi) as basal insulin analog formulations (33,34). This analog contains two types of modifications. The first is the addition of two basic residues at the C-terminus of the B chain (Arg^B31^-Arg^B32^), which shifts the isoelectric point to neutral pH, thereby making the protein molecule more soluble at pH 4 (as in an acidic pharmaceutical formulation) and yet insoluble at physiological pH (as in the subcutaneous depot). The second modification is replacement of Asn^A21^ by Gly, which is associated with enhanced chemical stability (33,34) due to protection from acid-catalyzed deamination of the A21 side chain (29,35).

We therefore undertook the chemical synthesis of the two component peptide chains of Se-glargine: A-chain variant [C6U, C11U, N21G] and a B-chain variant containing Arg^B31^-Arg^B32^ by SPPS; the latter B-chain analog was also be obtained from Lantus^®^ SoloStar^®^ pen by sulfitolysis (below).

Fluorenylmethyloxycarbonyl (Fmoc)-based SPPS was used to synthesize A chain[C6U, C11U, N21G], wherein Sec residues were inserted manually at positions 6 and 11. After cleavage and deprotection by standard conditions, the peptide was isolated in 9% yield (Fig. 2). The variant B chain was likewise synthesized and purified in 20% yield (Fig. 3). Oxidative sulfitolysis of a commercial formulation was also employed to obtain sulfonated glargine B chain (Fig. 4). With the Sec-modified Gly^A21^ A chain and sulfonated B chain in hand, we initiated the combination reaction, in which A chain[C6U, C11U, N21G] was dissolved in 0.1 M glycine buffer at pH 10.6 (at a final concentration of 0.6 mM), and the sulfonated B chain was added (final concentration of 0.5 mM). At this pH, precipitation of the B chain was observed presumably due to the Arg^B31^-Arg^B32^ extension, which shifts the isoelectric point of the peptide. The pH was adjusted to 11.2 (by addition of 0.5 M NaOH), which enabled the B chain to be completely dissolved, and then DTT was added at a stoichiometric concentration relative to the sulfonate groups present in the B chain (1 mM). The reaction was left at 4 ^o^C with exposure to air to permit oxidation (36). Aliquots were taken at successive time points, quenched with 0.1% trifluoroacetic acid (TFA) in water and left in the freezer until injected into the reversed-phase HPLC. Under these conditions, Se-glargine was observed after 5 h in small amount, and so the reaction was left overnight; aggregates of the variant B chain were also observable. After 24 h, Se-glargine was isolated (ca. 20% by HPLC integration; with 7% isolated yield, Fig. 5).

**Figure 2.**
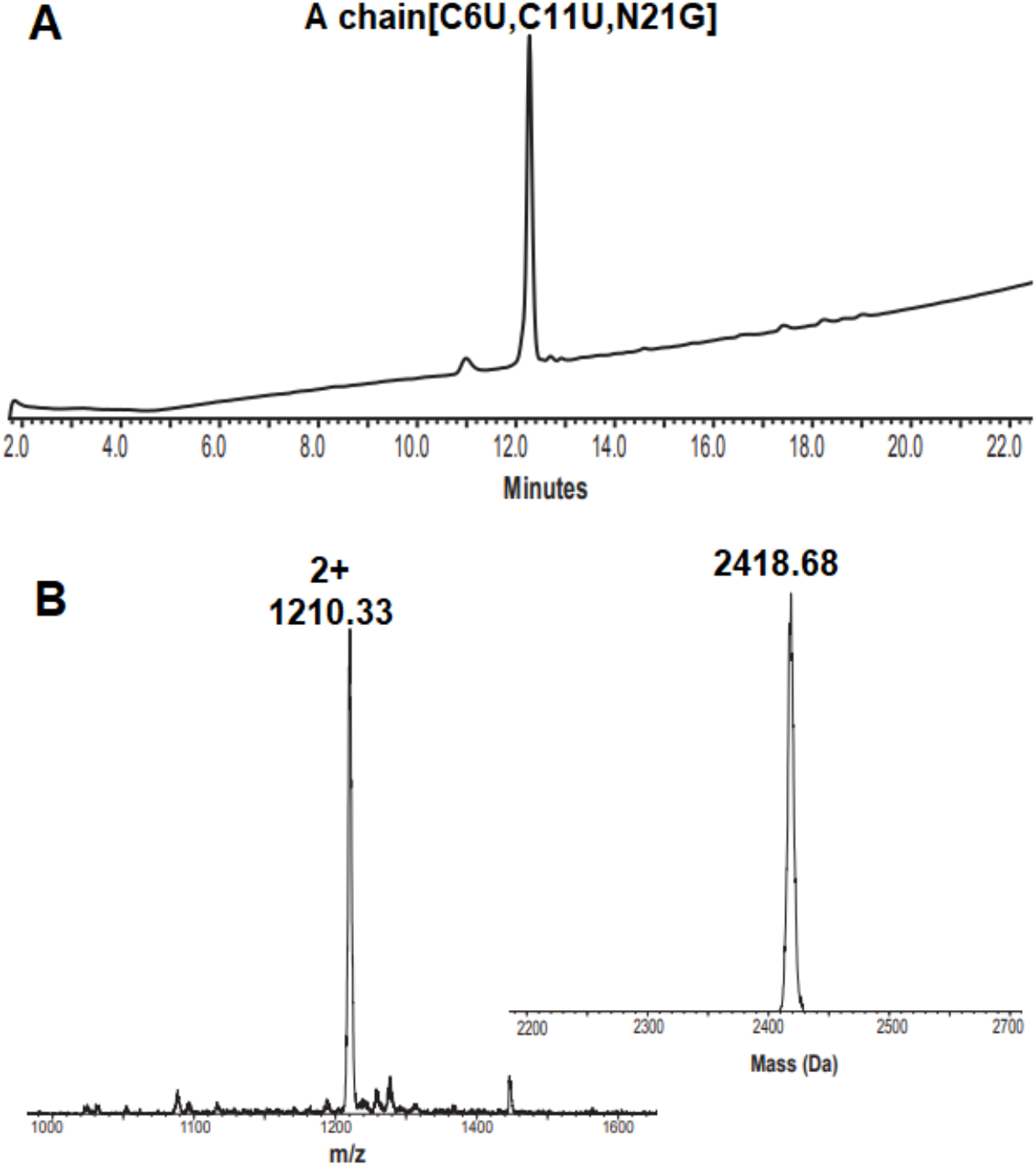
Characterization of A chain[C6U, C11U, N21G]. **A**. Analytical reversed-phase HPLC of the purified A chain[C6U, C11U, N21G], using column XBridge BEH300 C4 (3.5 μm, 130 Å, 4.6 × 150 mm, 5% B for 2 min then 5-70% B over 20 min, total 25 min); **B**. the corresponding ESI-MS, obs. 2418.68 Da, calc. 2418.44 Da, with two oxidation states, either two selenylsulfides or a diselenide and disulfide bonds.

**Figure 3.**
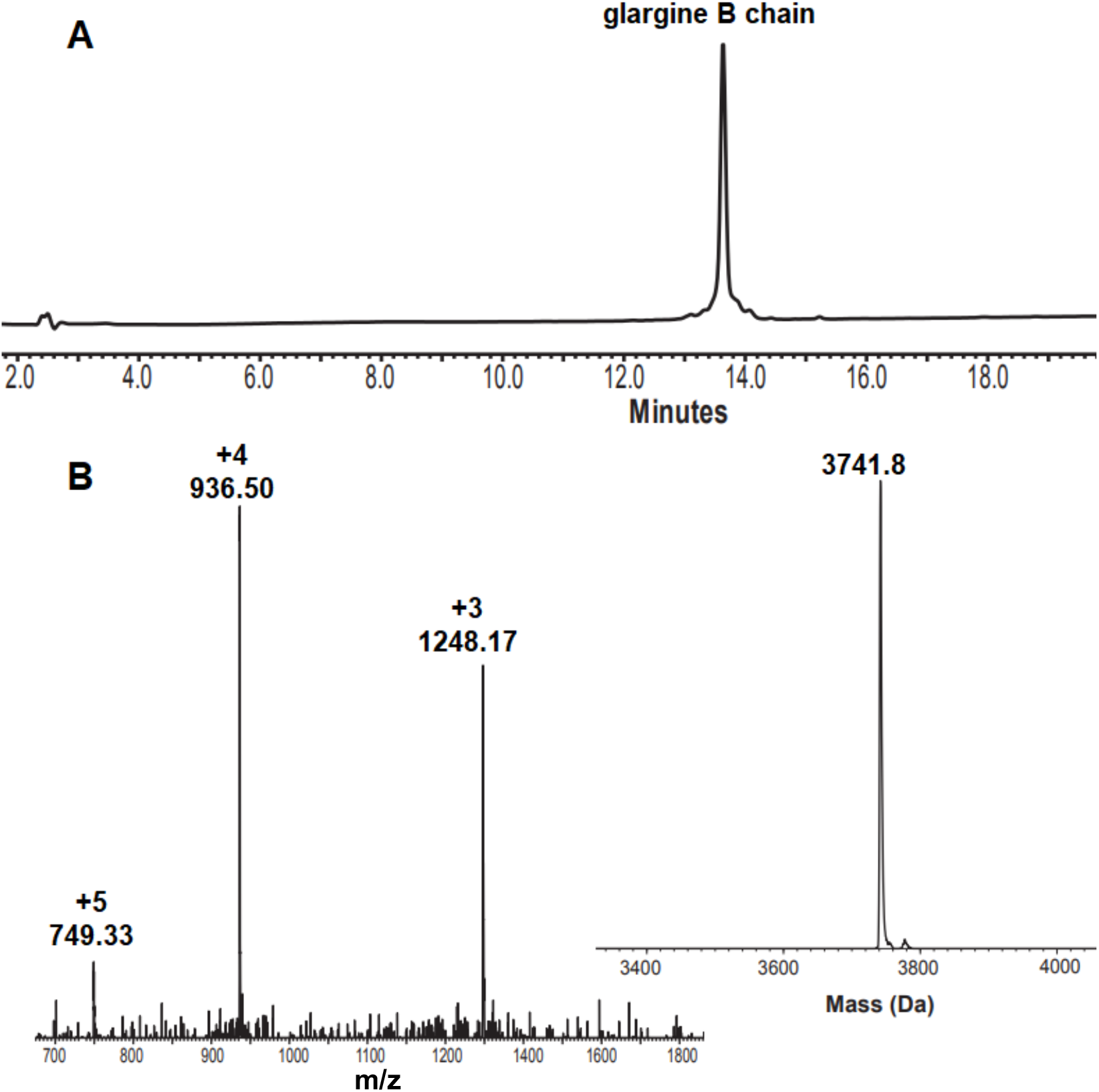
Characterization of glargine B chain. **A**. Analytical reversed-phase HPLC of the purified glargine B chain, column XBridge BEH300 C4 (3.5 μm, 130 Å, 4.6 × 150 mm, 5% B for 2 min then 5-70% B over 20 min, total 25min); **B**. the corresponding ESI-MS, obs. 3741.8 Da, calc. 3742.3 Da.

**Figure 4.**
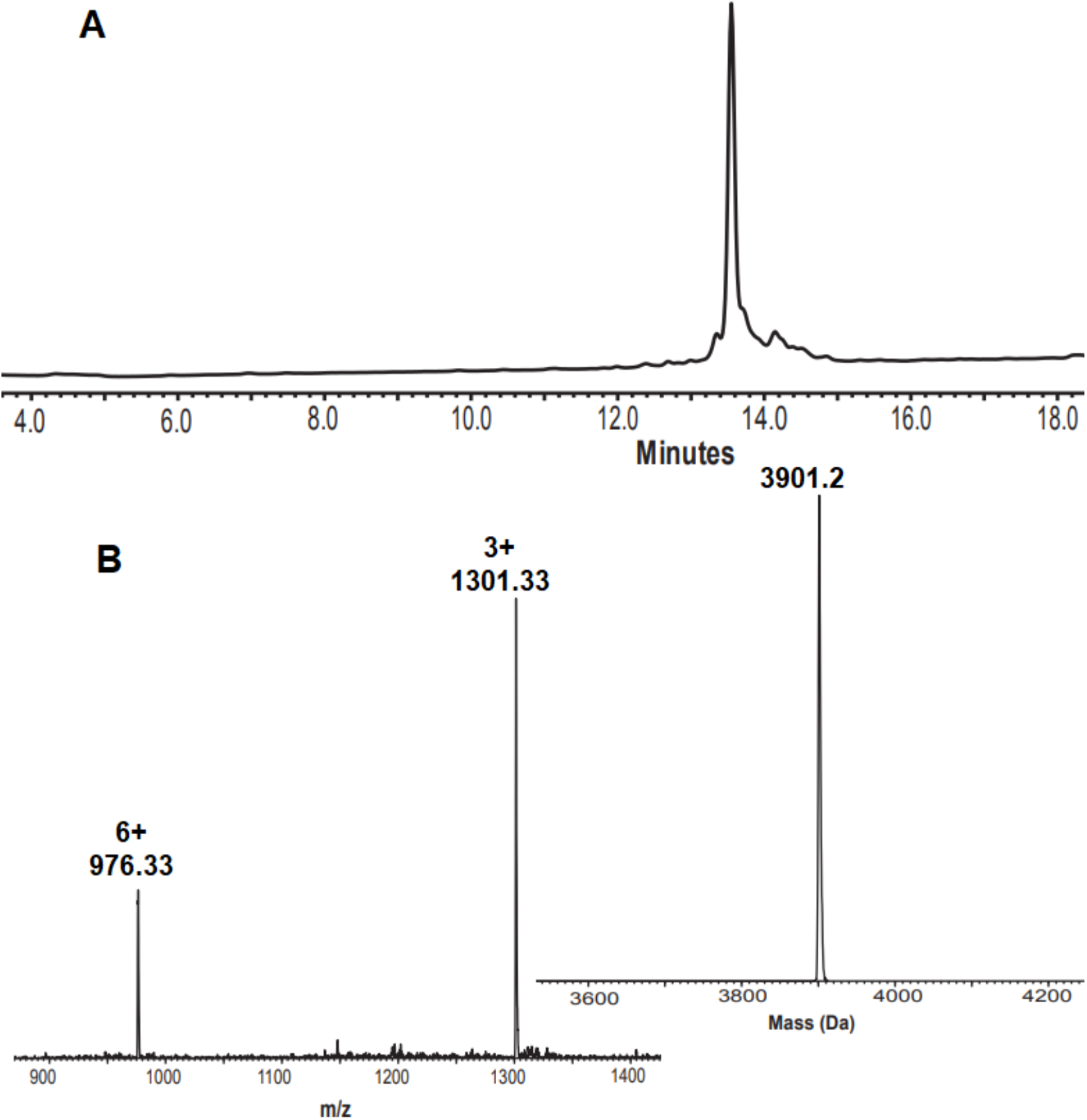
Analytical reversed-phase HPLC and ESI-MS of sulfonated synthetic glargine B chain. **A**. HPLC analysis of glargine B chain, glargine B chain(SSO_3_^2-^)_2_; **B**. The corresponding ESI-MS spectrum; obs. 3901.2 Da, calc. 3900.4 Da.

**Figure 5.**
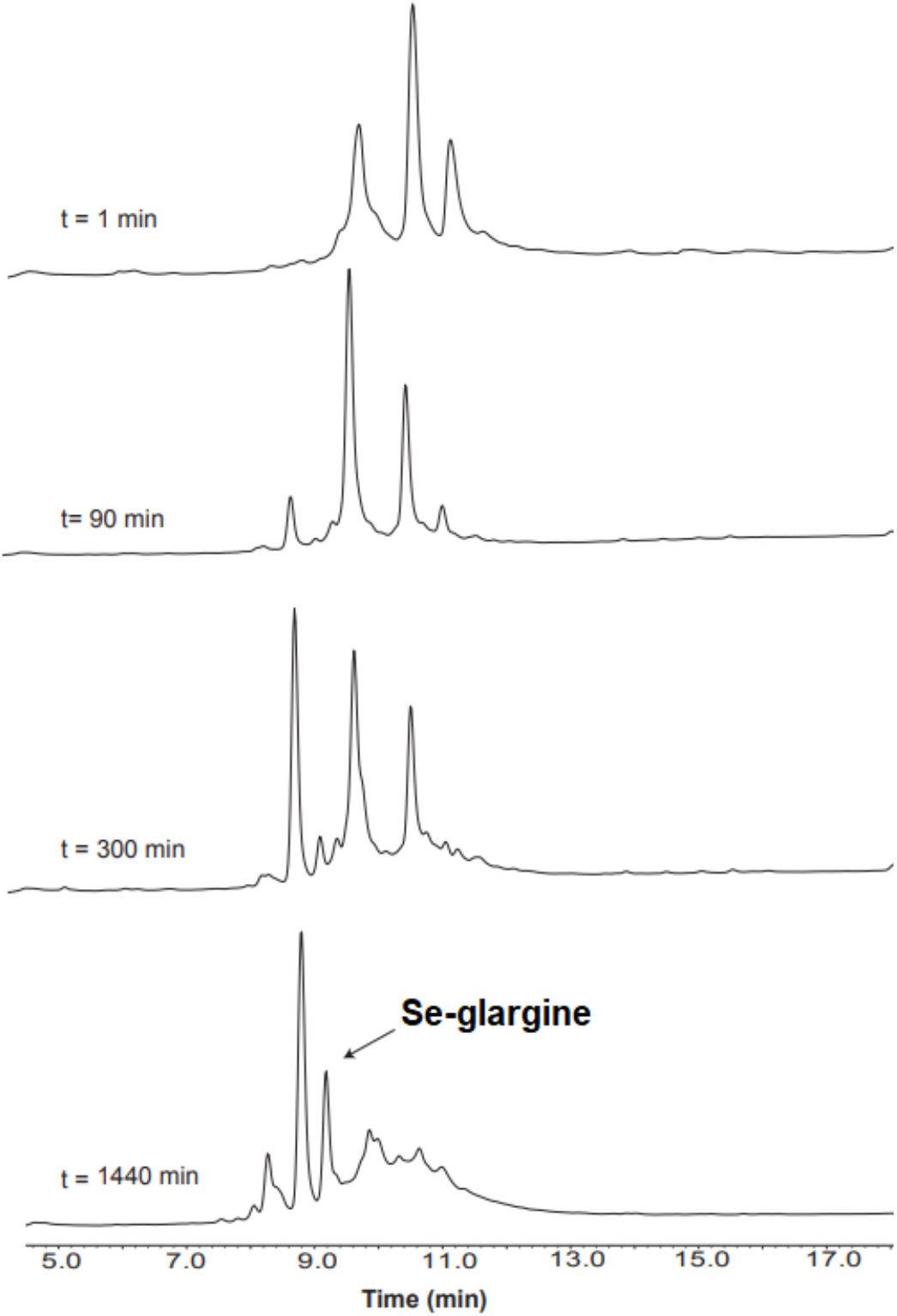
Analytical reversed-phase HPLC of the recombination reaction assay of Se-glargine using A chain[C6U, C11U, N21G] analog and sulfonated synthetic glargine B chain in a 0.1 M glycine buffer (pH∼11.2) at 4 ^o^C. Addition of DTT in quantity stoichiometric to the concentration of sulfonate groups initiated the reaction. Conditions: A chain[C6U, C11U, N21G] = 0.6 mM, [sulfonated B chain] = 0.5 mM, [DTT] = 1 mM. Se-glargine peak was observed after 5 h. Aggregation of the glargine B chain was observed at the end of the reaction (a broad peak R_t_ = 9-11).

The sulfonated glargine B chain was also isolated by oxidative sulfitolysis reaction (Fig. 6) applied on commercially available Lantus^®^ SoloStar^®^ pen (Sanofi), then purified by reversed-phase HPLC, and the same protocol for combination reaction was applied. Surprisingly, the yield of the combination reaction was significantly improved, and Se-glargine started to form after only 3 h; the reaction was completed after 18 h. The Se-glargine product was the major peak under these conditions, and A chain[C6U, C11U, N21G] was still present at the end of the reaction (which was taken in slight excess), while negligible amounts of the B chain were observed. Under these conditions, the Se-glargine was isolated in 20% yield (ca. 60% by HPLC integration; Fig. 7 and Discussion).

**Figure 6.**
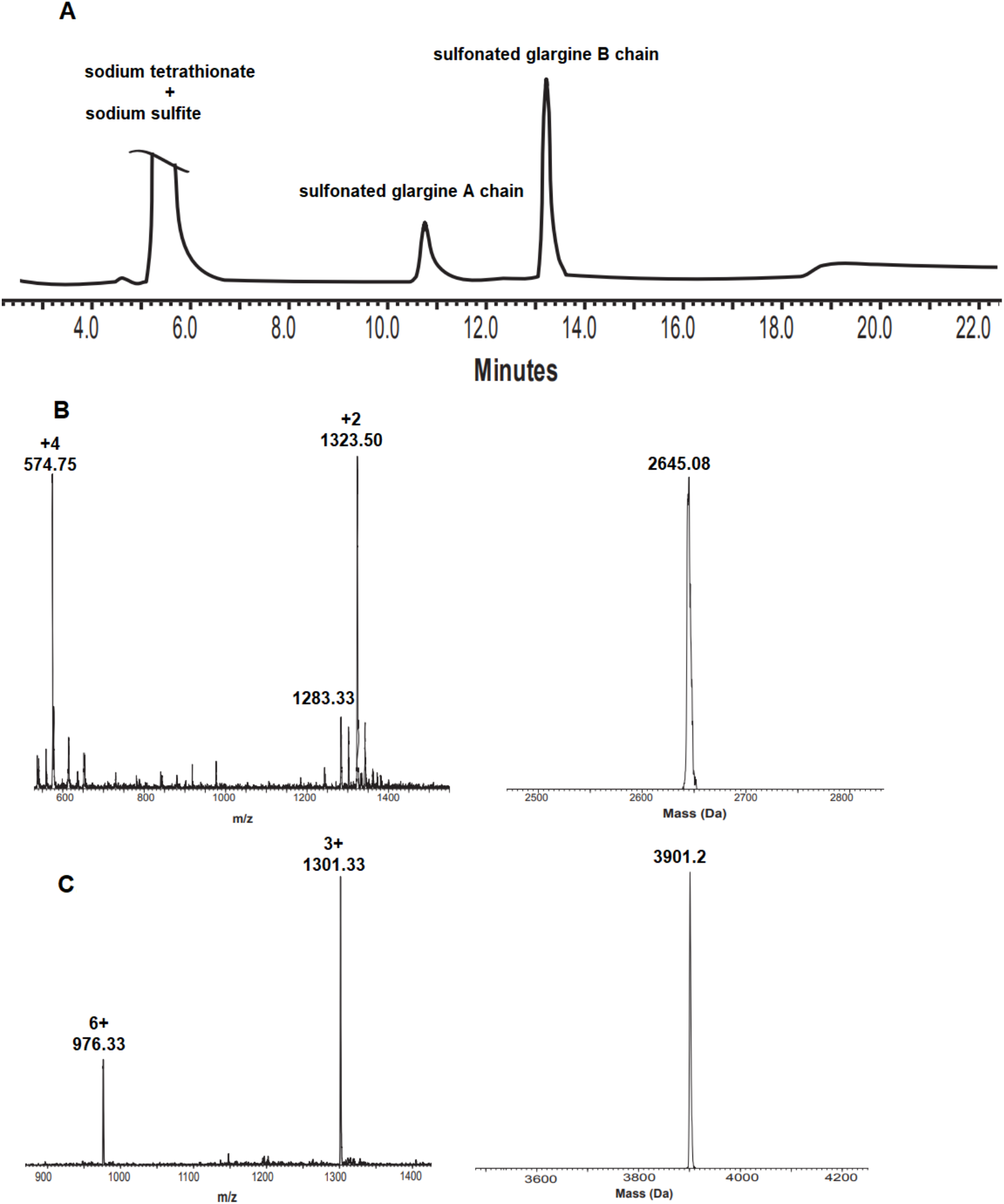
Analytical HPLC and ESI-MS of the sulfitolysis reaction of commercially available Lantus^®^ SoloStar^®^ pen (Sanofi). **a**. The sulfitolysis reaction after 2 h (see methods for detailed conditions). **b**. ESI-MS analysis of S-sulfonated A chain with 4 sulfonate groups: glargine A chain(SSO_3_^2-^)_4_ and its corresponding ESI-MS spectrum (obs. 2645.08 Da, calc. 2642.85 Da). **c**. ESI-MS spectrum of S-sulfonated glargine B chain; B chain(SSO_3_ ^2-^)_2_, (obs. 3901.2 Da, calc. 3900.44 Da).

**Figure 7.**
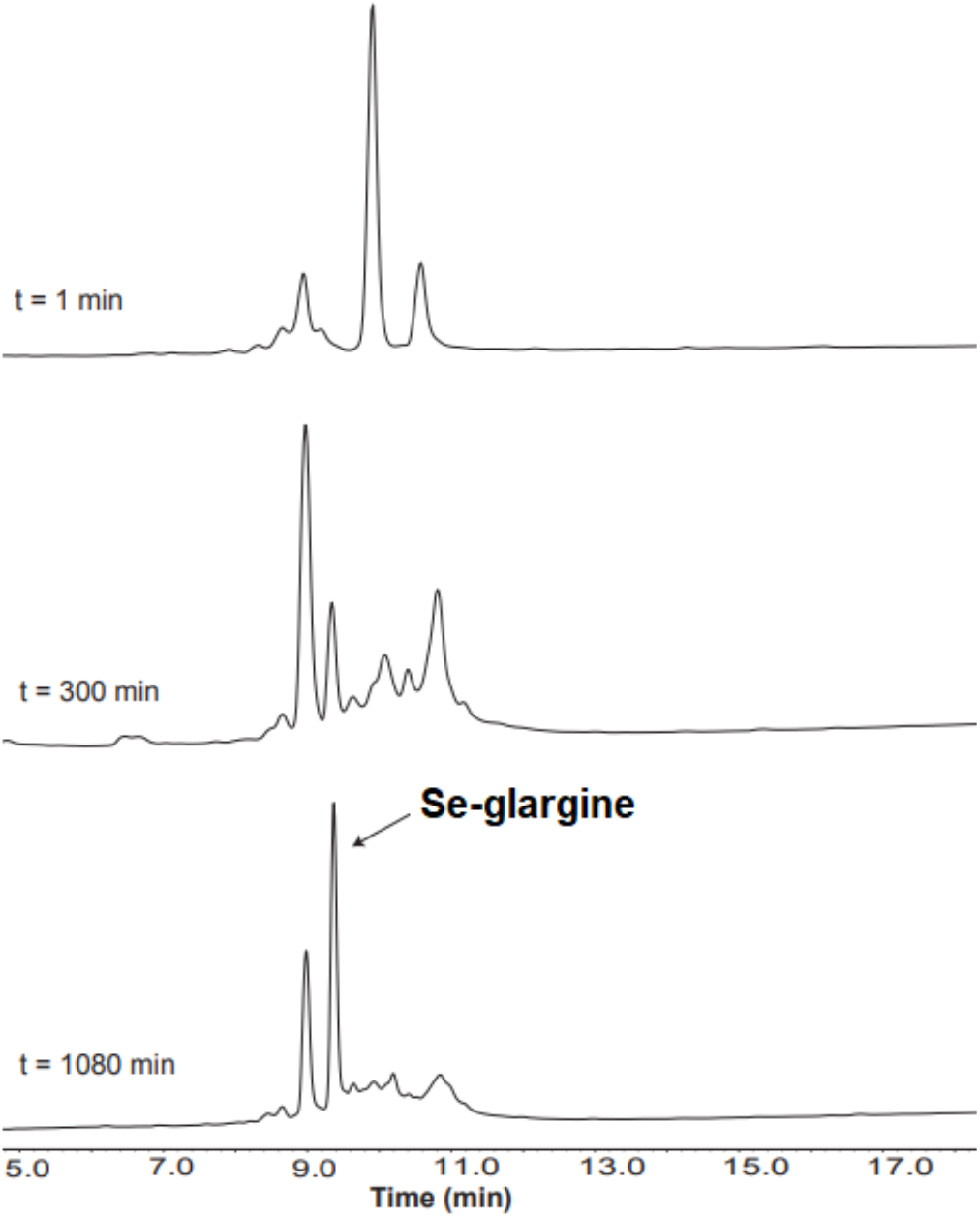
Analytical HPLC of combination assay for Se-glargine using chain A[C6U, C11U, N21G] analog with sulfonated B chain from Lantus^®^ Solostar^®^ pen in a 0.1 M glycine buffer (pH ∼ 11.2) at 4 ^o^C. Addition of DTT in quantity stoichiometric to the concentration of sulfonate groups initiated the reaction. Conditions: A chain[C6U, C11U, N21G] = 0.6 mM, [B chain] = 0.5 mM, [DTT] = 1 mM. The Se-glargine product was observed after 3 h in small amounts, and the reaction was completed after 18 h. Only minor aggregation of the glargine B chain was observed at the end of the reaction. The major peak is the Se-glargine product.

## Discussion

### Substitution of Cysteine for Selenocysteine

Selenocysteine (Sec, U), the 21^st^ encoded amino acid (10), is a near isostere of cysteine (Cys, C) that present unique chemical properties such as its low redox potential (E_0_ = -386 mV) and p*K*_a_ (∼5.2) (17) compared to Cys (12). Therefore, at physiological pH, the Sec side chain is predominantly deprotonated (selenolate), whereas the most Cys side chains remain protonated. These characteristics have been exploited in the field of protein folding *in vitro* (13-16,19-21).

Correct protein folding is critical for protein function and occurs spontaneously under suitable *in vivo* conditions, however, the *in vitro* folding process can be challenging in the case of Cys-rich proteins, such as insulin, due to the formation of non-native disulfide bonds and the formation of trapped intermediates, thus affecting the yield of properly folded proteins. Furthermore, Cys-to-Sec substitution has been demonstrated to enhance the oxidative folding of Cys-rich proteins *in vitro* without affecting the native conformations or biological activities of the target proteins (13,37).

In our previous paper, we observed that substitution of a single disulfide pair (at positions 6 and 11 in A chain) with Sec markedly enhanced chain-combination rate and yield while maintaining an indistinguishable biological activity from the wild-type, while maintaining the native like structure of insulin (4). Furthermore, the Se-insulin demonstrated enhanced stability, which was attributed to the enlarged atomic radii of the Se atoms (in Sec) in comparison to the S (in Cys), that afforded a better accommodation of the diselenide bond in the hydrophobic core without steric clash (not related to redox). This outcome was supported by detailed 2D-NMR and X-ray crystallographic studies (4).

### Challenges in chain combination of glargine

The Arg^B31^-Arg^B32^ extension in the glargine B chain shifts the isoelectric point of the chain and its pH-dependent solubility making the combination with the A chain more challenging than with regular B chain. (33,34). In fact, the glargine B chain is soluble in slightly more basic conditions compared to regular B chain (pH 11.2 *vs* 10.6), and this small change in the pH during the reaction can cause insolubility of the B chain and its aggregations. The use of glargine B chain isolated by oxidative sulfitolysis reaction applied on commercially available Lantus^®^ SoloStar^®^ pen (Sanofi) enhanced the yield (by a three-fold) and rate of the combination reaction, in which only negligible amounts of the B chain were observed at the end of the reaction. The observed improvement of the combination reaction is probably due to the presence of polysorbate 20, a surfactant presents in the composition of Lantus^®^ SoloStar^®^ pen, which is known to enhance the stability (38) that would remain present in low concentration after purification of the glargine B chain. We suggest that polysorbate 20 reduced formations of glargine B chain aggregates that are a competitive process to the combination reaction, and as such improve the yield of Se-glargine formation.

Se-glargine was isolated by HPLC (Fig. 8) and characterized by HR-MS (Fig. 9). In our companion study (2) we report that the native function (as probed in a cell-based model; 7) and structure (as probed by high-resolution NMR spectroscopy; 8) are maintained. Studies of shelf life and degradation rates are in progress in a formulation similar to those of Lantus or Toujeo (Sanofi). We anticipate that these foundational studies will lead to a basal insulin analog formulation of translational importance to global health in the face of an emerging pandemic of obesity-associated T2D (27,28).

**Figure 8.**
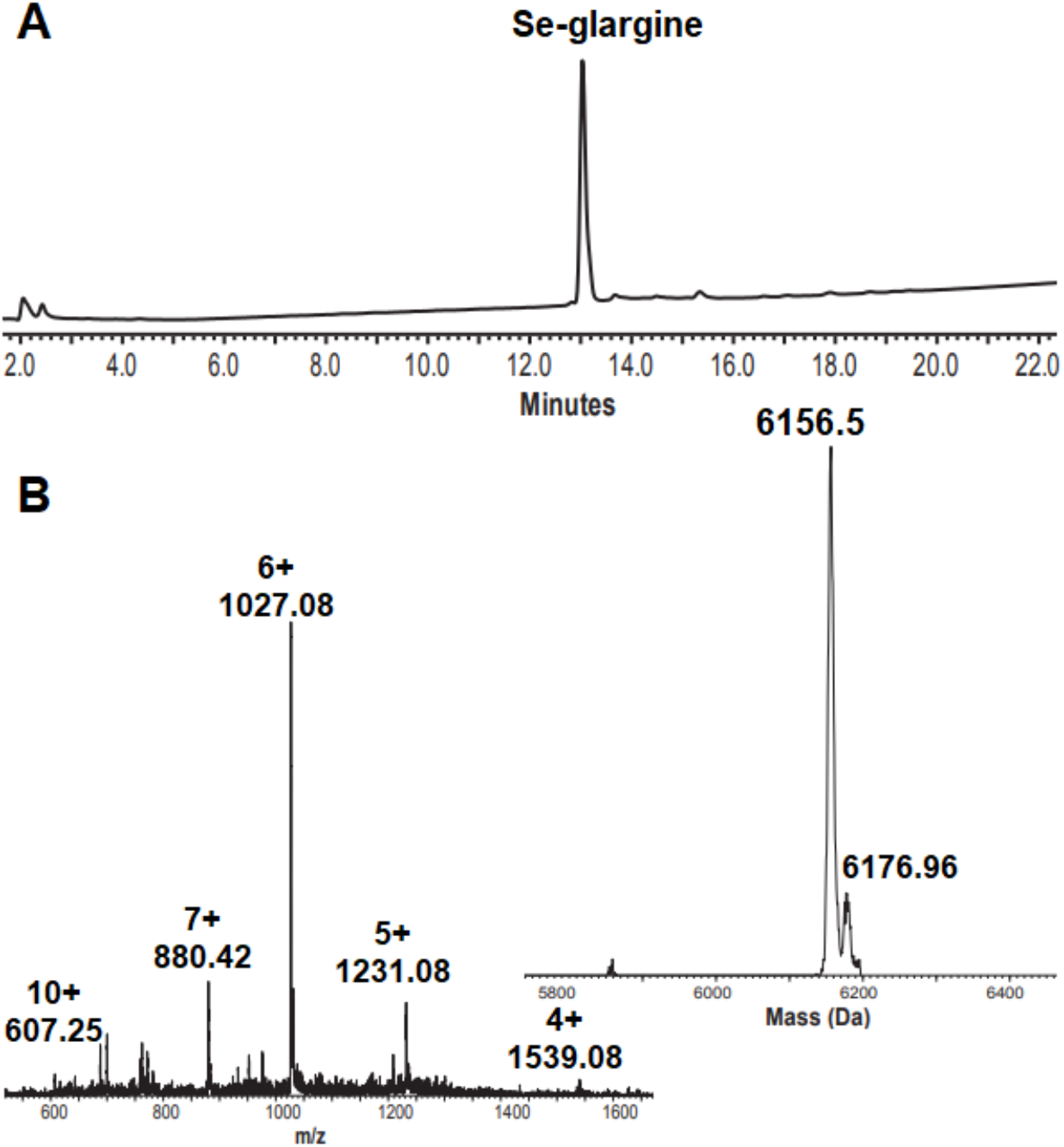
Characterization of Se-glargine. **A**. Analytical reversed-phase HPLC of the purified Se-glargine, XBridge BEH300 C4 column (3.5 μm, 130 Å, 4.6 × 150 mm, 10%-60%, in 20 min, total time 25 min); **B**. the corresponding ESI-MS, obs. 6156.5 Da, calc. 6156.75 Da.

**Figure 9.**
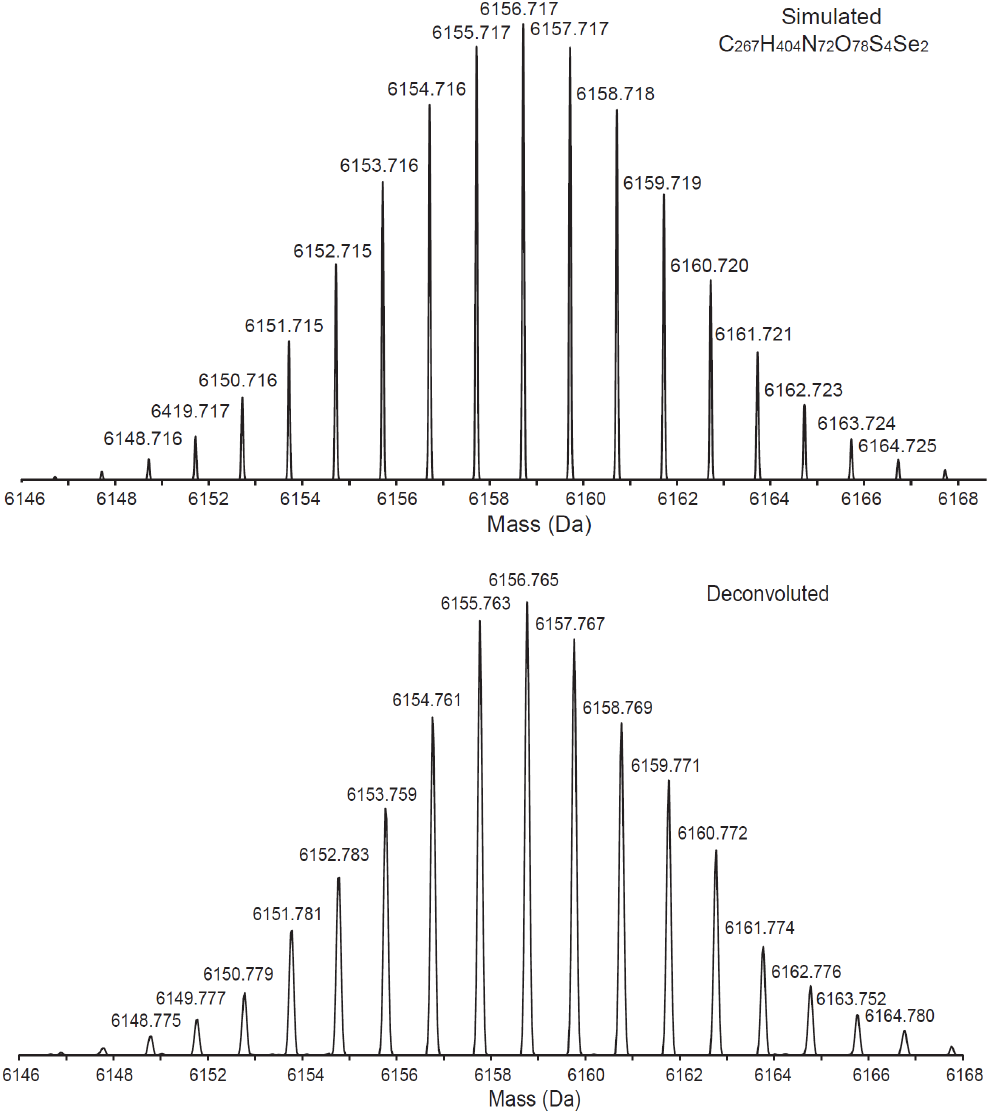
HR-MS analysis of Se-glargine. The simulated HR-MS of Se-glargine with chemical formula C_267_H_404_N_72_O_78_S_4_Se_2_ is shown (top); The deconvoluted HR-MS of Se-glargine (bottom).

## Experimental Procedures

### Materials

Buffers were prepared using MilliQ water (Millipore, Merck). Glycine hydrochloride buffer, purchased from Sigma-Aldrich (Rehovot, Israel), was used in all insulin recombination reactions. Similarly, NaS_4_O_6_·2H_2_O (sodium tetrathionate dihydrate), Urea, 2,2′-dithiobis(5-nitropyridine), ethanedithiol (EDT), *N*,*N′*-diisopropylcarbodiimide (DIC), thioanisol, triisopropylsilane (TIPS), *DL*-dithiothreitol (DTT), 2,2′-dithiobis(5-nitropyridine) (DTNP), ammonium bicarbonate (NH_4_HCO_3_) were purchased from Sigma-Aldrich (Rehovot, Israel) and Na_2_SO_3_ (sodium sulfite) from Merck. Ethylene diamine tetraacetic acid (EDTA) were obtained by J. T. Baker. Tris.HCl was purchased from Bio-Lab and NaH_2_PO_4_·H_2_O from JT Baker Chemicals. All standard Fmoc-amino acids were obtained from CS Bio Co. (Menlo Park, CA) or Matrix Innovation (Quebec City, Canada), with the following side chain protecting groups: Asn(Trt), Gln(Trt), Glu(OtBu), Gly(OtBu), Ile, Leu, Ser(tBu), Thr(tBu),Val, Cys(Trt), Tyr(tBu). 2-Chlorotrityl-resin (loading of 0.3-0.8 eq/g) was purchased from Chem-Impex International. HCTU and OxymaPure were purchased from Luxembourg Biotechnologies Ltd. (Rehovot, Israel). All solvents: *N*,*N*-dimethylformamide (DMF), dichloromethane (DCM), acetonitrile (ACN), *N*,*N*-diisopropylethyl amine (DIEA), trifluoroacetic acid (TFA) and piperidine (Pip) were purchased from Bio-Lab (Jerusalem, Israel) and were peptide synthesis, HPLC or ULC-grade. Pepsin endopeptidase from porcine gastric mucosa (≥250 units/mg) was purchased from Sigma-Aldrich (Rehovot, Israel). Lantus^®^ Solostar^®^ pens (Sanofi) were generously given by diabetes patient who no longer needed them. Fmoc-Sec(Mob)-OH synthesis was reported previously (39).

### High Performance Liquid Chromatography (HPLC)

Analytical reversed-phase HPLC (RP-HPLC) was performed on a Waters Alliance HPLC with 220 nm UV detection using a XBridge BEH300 C4 (3.5 μm, 130 Å, 4.6 × 150 mm). Semi-preparative RP-HPLC was performed on a Waters LCQ150 system using a XBridge C8 column (5 μm, 10 × 150 mm) or XBridge BEH C4 column (5 μm, 300 Å, 100 × 150 mm). Preparative RP-HPLC was performed on a XSelect C4 column (5 μm, 130 Å, 19× 250 mm) or XSelect C18 column (5 μm, 30 × 250 mm). The flow rates were 1 mL/min (analytical), 3.35 mL/min (semi-preparative), and 10-20 mL/min (preparative). Linear gradients of ACN (with 0.1 % TFA, eluent B) in water (with 0.1 % TFA, eluent A) were used for all systems to elute bound peptides. For purification of Se-glargine after combination, a different solvent system was used: a linear gradient of ACN (100%, eluent D) in 25 mM ammonium bicarbonate buffer (eluent C).

### Electrospray Ionization Mass Spectrometry (ESI-MS) and HR-MS

ESI-MS was performed on LCQ Fleet Ion Trap mass spectrometer (Thermo Scientific). Peptide masses were calculated from the experimental mass to charge (m/z) ratios from the observed multiply charged species of a peptide.

The HR-MS were recorded on a Q-ExactivePlus Orbitrap mass spectrometer (Thermo Scientific) with an ESI source and 140’000 FWHM, in a method with AGC target set to 1E^6^, and scan range was 400-2800 m/z. Deconvolution of the raw experimental MS data was performed with the help of MagTran v1.03 software.

### Peptide synthesis

#### Fmoc-SPPS synthesis of A chain[C6U, C11U, N21G] analog (21 AA)

The amino-acid sequence of the glargine-related variant A chain is G-I-V-E-Q-U-C-T-S-I-U-S-L-Y-Q-L-E-N-Y-C-G (pairwise Sec substitutions in *red*). This peptide (A chain[C6U, C11U, N21G]) was prepared by automatic peptide synthesizer (CS136XT, CS Bio Inc. CA) on a 0.25 mmol scale using 2-chlorotrityl-resin (loading of 0.3-0.8 eq/g). Fmoc amino acids (2 mmol in 5 mL DMF) activated with HCTU (2 mmol in 5 mL DMF) and DIEA (4 mmol in 5 mL DMF) for 5 min and allowed to couple for 25 min, with constant shaking. Fmoc-deprotection was carried out with 20% piperidine in DMF (2 × 5 min). Fmoc-Sec(Mob)-OH was manually coupled for two hours using DIC/OxymaPure activation method (2.9 equiv/3.0 equiv, respectively).

The peptide-resin was washed with DMF, DCM and dried under vacuum. The dried peptide-resin was cleaved in the presence of 2 equiv of DTNP, (40) using TFA: water: thioanisole: triisopropylsilane: ethanedithiol (94 : 1.5: 1.5: 1.5: 1.5) cocktail for 4 h. The cleavage mixture was filtered and TFA was evaporated with N_2_ bubbling to minimum volume, to which 8-fold volume of cold ether was added dropwise. The precipitated crude peptide was centrifuged (5000 rpm, 10 min), ether was removed and the crude peptide dissolved in ACN-water (1:1) containing 0.1% TFA and was further diluted to ∼25% ACN with water and lyophilized, giving 840 mg of crude material (including 240 mg of DTNP).

The crude peptide was dissolved in 25% ACN in water containing 0.1% TFA and purified by multiple injections of 50 mg each on prep RP-HPLC (XSelect C4 column, 5 μm, 19 × 250 mm) using a gradient of 25%-50% B over 50 min, to give 55 mg of chain A[C6U, C11U, N21G] analog (9% yield, based on resin loading), which was characterized by HPLC using XSelect C4 column (5 μm, 130 Å, 19× 250 mm), and a gradient of 5% B over 2 min then 5-70% B over 20 min and ESI-MS (Fig. 1).

#### Fmoc-SPPS synthesis of glargine chain B (32 AA)

The amino-acid sequence of the variant glargine B chain is F-V-N-Q-H-L-C-G-S-H-L-V-E-A-L-Y-L-V-C-G-E-R-G-F-F-Y-T-P-K-T-R-R. This 32-residue peptide was synthesized in a similar fashion as described for the A chain[C6U, C11U, N21G] analog. The peptide-resin was washed with DMF, DCM and dried under vacuum. The peptide-resin was washed with DMF, DCM and dried under vacuum. The dried peptide-resin was cleaved using TFA: water: thioanisole: triisopropylsilane: ethanedithiol (94 : 1.5: 1.5: 1.5: 1.5) cocktail for 4 h. The cleavage mixture was filtered and TFA was evaporated with N_2_ bubbling to minimum volume, to which 8-fold volume of cold ether was added dropwise. The precipitated crude peptide was centrifuged (5000 rpm, 10 min), ether was removed and the crude peptide dissolved in ACN-water (1:1) containing 0.1% TFA and was further diluted to ∼25% ACN with water and lyophilized, giving 805 mg of crude material. The crude peptide were dissolved in 25% ACN in water containing 0.1% TFA and purified by multiple injections of 100 mg each on prep RP-HPLC (XSelect C18 column, 5 μm, 30 × 250 mm) using a gradient of 30%-60% B over 50 min, to give 187 mg of chain B glargine (20% yield, based on resin loading), which was characterized by HPLC using ACQUITY HPLC XSelect C4 column (3.5 μm, 130 Å, 4.6 × 150 mm), and a gradient of 5% B over 2 min then 5-70% B over 20 min and ESI-MS (Fig. 2).

#### Oxidative sulfitolysis reaction applied on the synthetic glargine B chain

After purification, the glargine B chain was added to sulfitolysis buffer containing 0.2 M Na_2_SO_3_ and 0.2 M Na_2_S_4_O_6_ in a 20 mM Tris-HCl, 1 mM EDTA, 8 M urea (41). Peptide concentration was 10-15 mg/mL and the pH was adjusted to 6-7. The reaction was monitored by analytical HPLC (XSelect C4 column, 3.5 μm, 130 Å, 4.6 × 150 mm), and ESI-MS and was completed in 2 h. At the end of the reaction, ACN was added to enhance solubility of the sulfonated B chain before purification (25% of the final volume). Semi-preparative RP-HPLC XBridge Prep C8 column (5 μm, 10 × 150 mm) or Prep RP-HPLC (XSelect C4 column, 5 μm, 19 × 250 mm) were used to purify the products. B chain with two S-sulfonate groups was obtained, one group linked to each cysteine residue (Fig. 3).

#### Sulfitolysis of glargine in Lantus formulation

Insulin glargine from Lantus^®^ Solostar^®^ pen was taken (3.64 mg/mL, total volume 3 mL) and were added to 3 mL of 20 mM Tris-HCl, 1 mM EDTA, 8 M urea buffer (41). 0.2 M Na_2_SO_3_ and 0.2 M Na_2_S_4_O_6_ were added and the pH was adjusted to 6-7. The reaction was monitored by analytical HPLC (XSelect C4 column, 3.5 μm, 130 Å, 4.6×150 mm), and ESI-MS and was completed in 2 h. Prep RP-HPLC (XSelect C4 column, 5 μm, 19 × 250 mm) was used to purify the products. The glargine A chain with four S-sulfonate groups, and B chain with two S-sulfonate groups, were obtained, one group linked to each cysteine residue (Fig. 5).

#### Se-glargine combination reactions

The combination reaction for the Se-glargine preparation included A chain[C6U, C11U, N21G] with S-sulfonated glargine B chain (either synthetic or as isolated from a Lantus^®^ Solostar^®^ pen). A solution of A chain[C6U, C11U, N21G] analog (4 mg, final concentration in the combination experiment was 0.6 mM) was dissolved in a glycine buffer (0.1 M, pH 10.6. (36) The S-sulfonated glargine B chain (4.5 mg, final concentration in the combination experiment was 0.5 mM) was dissolved in the same buffer. At this pH, precipitation of the S-sulfonated glargine B chain was observed, 0.5 M NaOH was added until complete dissolution of the B chain glargine (∼200 μL). To initiate the reaction, the solution of S-sulfonated glargine B chain was added to the solution of A chain[C6U, C11U, N21G] analog, and DTT was added in quantity stoichiometric to the concentration of S-sulfonate groups (final concentration of 1 mM), and the reaction pH was 11.2. The reaction was carried out at 4 °C with open sample reaction to permit air oxidation. The progress of the reaction was followed by taking small aliquots (10 μL) of the reaction solution and quenched by addition of water containing 0.1% TFA (15 μL) and analyzed by injecting 20 μL of the sample onto the analytical HPLC (XSelect C4 column, 3.5 μm, 130 Å, 4.6 × 150 mm) using a gradient of 5% eluent B in eluent A for 1 min then 20-70% eluent B over 21 min, and detection at 220 nm (Fig. 4 and 6). At the end of the combination reaction, the Se-glargine was isolated through a semi-prep XBridge BEH C4 column (5 μm, 300Å, 100 × 150 mm) using a gradient of 5% ACN only (eluent D) in 25 mM ammonium bicarbonate buffer (eluent C) for 5 min then 25-40% eluent D over 45 min, and detection at 220 nm. Under these conditions the Se-glargine was obtained in 7% yield (∼0.5 mg determined by UV absorbance at 280 nm; when the synthetic glargine B chain was used) and 20% yield (1.5 mg determined by UV absorbance at 280 nm; when glargine B chain from Lantus^®^ Solostar^®^ pen was used) based on the amounts of the chains used (Figures 7 and 8).

## Abbreviations

Amino acids are designated by standard three-letter or one-letter code, with selenocysteine abbreviated as Sec or U. Residues within insulin are typically indicated by three letter code with chain and residue number in superscript; *e*.*g*., asparagine or glycine at position A21 of the A chain would be designated Asn^A21^ or Gly^A21^.

## Author Contributions

Peptide synthesis, chain combination, purification were performed by O.W.-K. with the advice of B.D., M.A.W. and N.M. The overall program of research was guided by M.A.W. and N.M. All authors contributed to preparation of the manuscript.

## Acknowledgements

We thank members of our respective laboratories for discussion. O.W-K was supported by the Kaete Klausner Ph.D. scholarship. N.M. acknowledges the financial support of the Israel Science Foundation (1388/22). M.A.W. acknowledges the support of the U.S. National Institutes of Health (NIH R01 DK04949) and the INCITE program of the Lilly Foundation at the Indiana University School of Medicine. B.D. and M.A.W. thank *emeritus* Prof. Stephen B. Kent (University of Chicago) for encouragement.

## FOOTNOTES

### Disclosure

O.W.-K., N.M. and M.A.W. hold a US patent number PCT29920-315514 entitled “stabilization of prandial or basal insulin analogues by an internal diselenide bridge” that covers this study.

^1^Unlike an internal diselenide bridge at position A6-A11, an external diselenide bridge at B7-A7 does not stabilize the insulin molecule as probed by chemical-denaturation studies (42). Because the thermodynamic stability of a protein reflects difference between the free energies of the bridged native state and bridged denatured state (as distinct from the reduced and unfolded state), any intrinsic physico-chemical differences in the redox properties between Cys and Sec (footnote 2) do not in themselves contribute to net free-energy differences (ΔΔG_u_). By contrast, differences in steric properties betweena disulfide bridge and diselenide bridge can in principle introduce favorable or unfavorable changes in core packing efficiency (42).

^2^The selenol group of Selenocysteine (Sec) has a lower pK_a_ (near 5.2) and lower reduction potential (E_0_= -388 mV) than does the thiol group of Cys; these differences rationalize its evolution within the active sites of seleno-enzymes and under its use as a probe in redox chemistry.

